# BicGO: a new biclustering algorithm based on global optimization

**DOI:** 10.1101/406769

**Authors:** Guojun Li, Zhengchang Su

**Affiliations:** School of Mathematics, Shandong University, Jinan 250100, China; Department of Bioinformatics and Genetics, University of North Carolina at Charlotte, NC28262, USA

## Abstract

Recognizing complicated biclusters submerged in large scale datasets (matrix) has been being a highly challenging problem. We introduce a biclustering algorithm BicGO consisting of two separate strategies which can be selectively used by users. The BicGO which was developed based on global optimization can be implemented by iteratively answering if a real number belongs to a given interval. Tested on various simulated datasets in which most complicated and most general trend-preserved biclusters were submerged, BicGO almost always extracted all the actual bicluters with accuracy close to 100%, while on real datasets, it also achieved an incredible superiority over all the salient tools compared in this article. As far as we know, the BicGO is the first tool capable of identifying any complicated (e.g., constant, shift, scale, shift-scale, order-preserved, trend-preserved, etc), any shapes (narrow or broad) of biclusters with overlaps allowed. In addition, it is also highly parsimonious in the usage of computing resources. The BicGO is available at https://www.dropbox.com/s/hsj3j96rekoks5n/BicGO.zip?dl=0 for free download.

## Introduction

Biclustering is to find sub-matrices with specific row patterns under certain columns in a numeric matrix. Biclustering a matrix is often desired in many fields of data analysis. For example, biologists would like to bicluster a gene expression matrix with rows corresponding to genes and columns to conditions/samples, under which the mRNA expression levels of genes are measured using various high-throughput methods such as DNA microarrays and more recently RNA-seq. The goal of biclustering such gene expression matrices is to find groups of genes with similar mRNA expression patterns under different groups of conditions [1, 2]. Comparing the gene expression patterns under different samples such as cancerous tissues at different developmental stages, one may derive information about genes associated with specific biochemical pathways dysregulated during the development of cancer. The biclustering problem can be traced back to Morgan et al. [3, 4] and Hartigan [5, 6] who attempted to partition a numerical matrix into submatrices whose values were as similar as possible. Since Cheng and Church [1, 2] firstly introduced a biclustering algorithm for gene expression data analyses in which mean squared residue was used as a measure, lots of biclustering algorithms [7-14] have been developed based on the measure. It was lately recognized that the mean squared residue measure is not adequate in application of detecting transcriptionally co-regulated genes [15-17]. Aguilar-Ruiz [18, 19] proposed more general measures, including shifting, scaling and their combination, to characterize co-regulated genes, though discovering these types of biclusters could be highly challenging. To tackle these challenges, one of us [22] introduced QUBIC, a qualitative biclustering algorithm that solved the more general problem to some extent, but still has limitations (see the counterexample of QUBIC in Supplementary Materials).

More recently, it is realized that the most biologically relevant gene expression patterns tend to be trend-preserved. Two genes’ expression is said to be trend-preserved under certain conditions if and only if their corresponding vectors in the matrix are either order-preserved or order-reversed, where two vectors *x* and *y* are said to be order-preserved if and only if any two corresponding components have the same rank (with respect to the numerical value) in their respective vectors, and order-reversed if and only if *x* and -*y* are order-preserved. A bicluster is said to be trend-preserved if and only if any pair of rows within the bicluster are trend-preserved. For general applicability, the entries within a trend-preserved bicluster are allowed to be same. Trend-preserved biclusters are most biologically meaningful structures in gene expression data because genes that are co-regulated may have quite different expression levels under different conditions, but often show trend-preserved expression patterns. Obviously, the widely studied biclusters with values being constant [20, 21], shifting [18, 19], scaling [18, 19], or shifting-scaling [18, 19], or order-preserved [15], etc. are all trend-preserved (see Supplementary Materials for details). Unfortunately, the problem of discovering trend-preserved biclusters is computationally intractable as even the simplest case with constant values is NP-hard [20, 21]. Wang et al. [22] presented a biclustering algorithm UniBic for identifying trend-preserved biclusters by applying the longest common subsequence (LCS) algorithm [23] to a new index matrix derived from the original matrix. The UniBic improved QUBIC to some extent, but it has to be collapsed when the biclusters to be detected are relatively narrow, i.e., very few columns and great many rows. Methodologically, locating seeds in UniBic procedure is not able to guarantee a global optimum solution (see Supplementary Materials for a counterexample). Very recently, when preparing the manuscript, we found a new biclustering algorithm EBIC introduced by Orzechowski, et al. [24] developed based on evolutionary computation to make up for the shortages inherited from UniBic [22]. However, the motivation in conceiving EBIC was not built on a right root. Actually, adding a new column into the current trend-preserved bicluster *B(I, J)* will generate *|J|+1* trend-preserved biclusters *B*(*I’, J’*) with |*I’*| approximating *|I|*/(*|J|+1*) instead of *|I|/2* (see Method). In addition, all the existing biclustering algorithms but EBIC [24] and BicPAMS [25] tend to be collapsed when the biclusters are relative narrow, i.e., with few columns and great many rows, because they were all developed based on maximal row-sequential patterns. BicPAMS [25] was developed specifically for narrow biclusters, but it performs poorer than EBIC [24], and even collapsed when data matrix is relatively broad due to its brute enumeration nature.

Motivated from the observation that there is a column permutation in a trend-preserved bicluster such that each permutated row is monotonic, i.e., increased or decreased (not necessarily strict), we introduced a new biclustering algorithm BicGO which substantially revolutionized the traditional approaches in all aspects. The BicGO consists of two separate strategies: column-based and row-based, which can be selectively used by users. The surprised is that the first strategy can be implemented by iteratively answering if a real number belongs to a given interval, while the second strategy can be implemented by iteratively finding a longest path in specific directed acyclic graphs (see Mehtod). Not only can the BicGO be implemented so easily, but also it is unprecedentedly powerful for identification of any complicated and any shapes of biclusters with high tolerance to overlaps. Although both strategies can be used universally, the first is more favorite for narrow and the second for broad.

Tested on various simulated datasets in which most complicated and most general trend-preserved biclusters were implanted and totally submerged, BicGO almost always extracted all the actual bicluters with unprecedented accuracy (always close to 100%), while on real datasets, it also achieved an incredible superiority over all the salient biclustering tools compared in this article. As far as we know, the BicGO is the first tool capable of identifying any complicated (e.g., constant, shift, scale, shift-scale, order-preserved, trend-preserved, etc) and any shapes (narrow or broad) of biclusters with overlaps allowed. In addition, it is also highly parsimonious in the usage of computing resources.

## Methods

As mentioned in previous literatures, it is highly challenging to recognize all the complicated biclusters submerged in various noisy data matrices, e.g., gene expression microarray datasets. However, we observed that the most complicated biclusters, e.g., trend-preserved ones no matter narrow or broad, could be recognized from a messy matrix as easily as constant biclusters are recognized from a binary matrix. We thus developed a new biclustering algorithm BicGO based on global optimization that consists of two separate strategies, column-based and row-based, which can be selectively used by users. The column-based strategy was implemented by iteratively answering if a real number belongs to a given interval, while the row-based strategy was implemented by iteratively finding the longest path in a specific directed acyclic graph constructed using a pair of sequences. In addition, we can narrow down the search space for each strategy based on the significance of biclusters to be identified.

### The BicGO algorithm

In the real world, the biclusters to be detected are usually very narrow of few columns and great many rows. However, all the existing tools but BicPAMS [30] and EBIC [24] tend to collapse in this situation. It is thus imperative to develop a new biclustering algorithm which is more powerful for recognizing not only broader but also narrower biclusters.

### Column-based strategy

The key step in this strategy is to locate seeds based on columns. In terms of definition, each pair of columns form an *<u>m×2</u>* trend-preserved bicluster, and thus a nontrivial seed should anchor at least three columns. Intuitively, if a triple of columns all pass through an actual bicluster, then they can be used to anchor a genuine seed. Obviously, all the actual biclusters can be easily obtained by growing their respective genuine needs. Thus, our goal is to first find out all the most significant seeds each is a trend-preserved *h×3* submatrix, where *h* is called the height of the seed. Certainly, the higher the height of a seed, the higher the probability of the seed being genuine. At first glance, it seems not easy to identify the most significant seed (trend-preserved *h×3* bicluster) from a triple of columns. In fact, this is a combinatorial optimization problem with itself being quite interesting. Motivated from the observation mentioned in introduction, we can model this seed optimization problem as to repeatedly deciding whether or not a real number falls into a given interval of type [*a, b*], (*-∞, b*] or [*a, +∞*). Given two columns *j*_*1*_ and *j*_*2*_, we greedily add a new column *j’* to get the most significant candidate seed, i.e., a trend-preserved *h×3* bicluster with the height *h* maximized, anchored on the three columns {*j*_*1*_, *j*_*2*_, *j’*}. We now elaborate how to locate the most significant candidate seeds determined by each pair of columns.

For each pair of columns (*j*_*1*_, *j*_*2*_), we denote by *A(I, J)* a *|I|×|J|* matrix, where *I = {1, 2, …, m}* and *J = {j*_*1*_, *j*_*2*_*}*. We call row *i* increased if 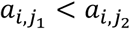, decreased if 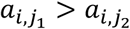 and identical if 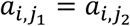 for *i* = 1, …, *m*. Considering a new column *j’*, we form three matrices *A(I, J*_*i*_*), i = 0, 1, 2,* with *J*_*0*_ *= {j’, j*_*1*_, *j*_*2*_*}, J*_*1*_ *= {j*_*1*_, *j’, j*_*2*_*}* and *J*_*2*_ *= {j*_*1*_, *j*_*2*_, *j’}*. Then the most significant candidate seed anchored on the three columns *j*_*1*_, *j*_*2*_, *j’* can be obtained from one of *A (I, J*_*0*_*), A (I, J*_*1*_) and *A (I, J*_*2*_) by deleting the rows violating monotonicity.

*Violation decision:* In *A (I, J*_*0*_), a row *i* violates monotonicity if and only if either 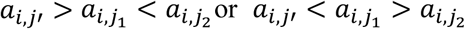. Let *A(I*_*0*_, *J*_*0*_) be the submatrix obtained from *A(I, J*_*0*_) by deleting the rows violating monotonicity. In *A (I, J*_*1*_), a row *i* violates monotonicity if and only if *a*_*i,j*′_ does not belong to the closed interval with boundaries 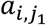 and 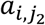. Let *A(I*_*1*_, *J*_*1*_) be the submatrix obtained from *A(I, J*_*1*_) by deleting the rows violating monotonicity. In *A(I, J*_*2*_), a row *i* violates monotonicity if and only if either 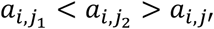 or 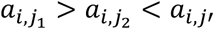. Let *A(I*_*2*_, *J*_*2*_) be the submatrix obtained from *A(I, J*_*2*_) by deleting the rows violating monotonicity. Denote

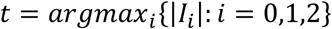

Then *A*(*I*_*t*_, *J*_*t*_) is the most significant candidate seed anchored on the given three columns *j*_*1*_, *j*_*2*_ and *j’.* Greedily enumerating *j’*, we obtain a most significant candidate seed determined by the columns *j*_*1*_ and *j*_*2*_. Repeat the procedure over all the column pairs, we thus obtain all the significant candidate seeds each is determined by a pair of columns, and then sort them in a candidate list *L* in an order decreasing in height. A genuine seed can be potentially grown into an actual bicluster by greedily adding a new column one at a time in a same way until the current bicluster goes down in significance, while fake ones will fall into disuse. To speed up the algorithm, the columns of the original data matrix can be partitioned into *k-1* subsets, where the integer k is the minimum number of columns the actual biclusters should have.

### Pseudo-codes of column-based strategy

#### Step 1. Partition of the columns

*We* calculate *k* based on the data matrix [15], and equally divide the columns into *k-1* subsets.

#### Step 2. Generation of candidate seeds

We calculate all the significant candidate seeds determined by each column pair from each column subset obtained in Step 1, and sort them in a candidate list *L* in an order descending in height.

#### Step 3. Growth without noise

We start with the first candidate *S* = (*I, J*), *J* = (*j*_1_, *j*_2_, *j*_3_) in *L*. Considering a new column *j’*, we form four matrixes *A*(*I, J*_*i*_), *i = 0, 1, 2, 3*, where *J*_0_ = {*j*^′^, *j*_1_, *j*_2_, *j*_3_}, *J*_1_ = {*j*_1_, *j*^′^, *j*_2_, *j*_3_}, *J*_2_ = {*j*_1_, *j*_2_, *j*^′^, *j*_3_ } *and J*_3_ = {*j*_1_, *j*_2_, *j*_3_, *j*^′^ }. Let *A*(*I*_*i*_, *J*_*i*_), *i = 0, 1, 2, 3,* be *|I*_*i*_*|×4* trend-preserved bicluster obtained from *A(I, J*_*i*_*), i = 0, 1, 2, 3,* by deleting the rows violating monotonicity, and denote

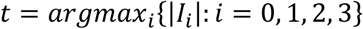

Then *A(I*_*t*_, *J*_*t*_) is the most significant *|I*_*t*_*|×4* trend-preserved bicluster determined by the four columns *j*_*1*_, *j*_*2*_, *j*_*3*_ and *j’.* Greedily enumerating *j’*, we obtain a most significant four-column trend-preserved bicluster, still denoted by *A*(*I*_*t*_, *J*_*t*_), which is determined by the columns *j*_*1*_ and *j*_*2*_. Repeating the procedure on *A*(*I*_*t*_, *J*_*t*_) until the current bicluster goes down in significance, we then obtain a so-called core bicluster, with which we go to the next step.

#### Step 4. Growth with noise

We extend the core bicluster obtained in Step 3 by adding as many rows of length at least *(1-c)|J|* as possible where *c* is an error rate (default: *0.01*). We remove the candidate seeds from the list *L* if their root column pairs are included in the discovered bicluster. Repeat step 3 until *L* has been exhausted or the required number of biclusters have been obtained.

#### Step 5. Output

Output all the biclusters obtained in Step 4.

### Row-based strategy

This strategy is more favorite for identifying broader biclusters submerged in various messy data matrices. Intuitively, if a pair of rows passing through an actual trend-preserved bicluster, then they can be used to grasp a genuine seed, the most significant trend-preserved *2×l* submatrix, where *l* is called the length of the seed. The genuine seed can grow to the full-sized bicluster to be detected, while fake one will fail to do so. Thus, our goal is to first find out at least one genuine seed for each actual bicluster. Obviously, the longer the length of a seed, the higher probability of the seed being genuine. We thus model this seed optimization problem as to finding a longest path in a specific directed acyclic graph constructed using a pair of sequences. Specifically, given a pair of rows *r*_*i*_ and *r*_*j*_, we construct a graph *G*_*ij*_ using the columns as the nodes and making an edge from column s to column t (*s ≠ t*) if and only if 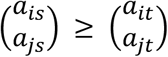, where “=“ holds if and only if both *a*_*is*_ = *a*_*it*_ and *a*_*ijs*_ = *a*_*jt*_. Therefore, an edge *(s, t)* is bidirectional if and only if 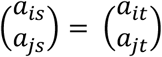. Then the graph *G*_*ij*_ has the following properties:

1. Let 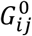 be a graph induced from all the bidirectional edges of *G*_*ij*_, then each component of 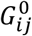 is a clique (there is a bidirectional edge between each pair of nodes in the component) of at least two nodes;
2. Let 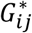 be a directed graph obtained from *G*_*ij*_ by contracting each clique into a node, then 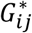 is acyclic;
3. The longest directed path in *G*_*ij*_ can be trivially obtained from 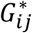 by blooming the cliques along the directed paths in 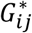 (see Supplementary Materials for details);
4. The longest directed path in *G*_*ij*_ corresponds to the longest seed, a *2×l* trend-preserved bicluster, determined by rows *r*_*i*_ and *r*_*j*_.

Based on pigeonhole principle, we partition the rows of the data matrix into *k-1* subsets to speed up the algorithm without loss of optimality, where *k* is the minimum number of rows actual biclusters should have.

### Pseudo-codes of row-based strategy

#### Step 1. Partition of the rows

We calculate an integer *k* using the mothed developed by Ben-Dor et al. [15] (see Supplementary Materials for details), and equally divide the rows into *k-1* subsets.

#### Step 2. Generation of seed candidates

We find all the candidate seeds by finding a longest path in each graph *G*_*ij*_ constructed using the pair of rows *r*_*i*_ and *r*_*j*_ from each subset obtained in Step 1, and then sort them in an order decreasing in length forming a candidate seed list *L*.

#### Step 3. Growth without noise

We start with the first seed in *L*, and greedily add a new row or its opposite into the current bicluster (seed) by finding a longest path in graph *G* constructed using the seed and the new row in a same way as *G*_*ij*_ was constructed. Repeating the step until the current bicluster goes down in significance, we then go to next step.

#### Step 4. Growth with noise

We start with the core bicluster obtained in step 3, and extend it as follows:

*Step 4.1.* Add new columns one at a time with an error rate *α* (default: 0.33) until none is available (see Supplementary Materials for details).

*Step 4.2.* Add new rows one at a time with an error rate *β* (default: 0.33) until none is available.

#### Step 5. Reset the seed list

We remove from the list *L* those with their row pairs passing through the bicluster obtained in Step 4, and then repeat Step 3 until *L* has been exhausted or the required number of biclusters have been obtained.

#### Step 6. Output

Output all the biclusters obtained in Step 4.

### Evaluation Criterion

Since there is no golden benchmark real dataset to validate the accuracy of a biclustering algorithm, it is a common practice to test it on simulated datasets using two measures, recovery and relevance scores, introduced by Prelić et al. [16], based on the match scores (Jaccard coefficients) [16] between the predicted and actual biclusters. Concretely, for two biclusters b_1_ and b_2_, the match score between them is defined as the ratio between the number of genes in their intersection and the number of genes in their union, i.e., *ms(b*_*1*_, *b*_*2*_*) = |b*_*1*_∩*b*_*2*_ *|/ |b*_*1*_∪*b*_*2*_ *|*, which measures the similarity between the two biclusters. For two sets of biclusters *M*_*1*_ and *M*_*2*_, the match score between them is defined as [16]

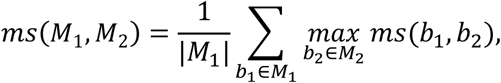

which measures the average similarity between biclusters in *M*_*1*_ and *M*_*2*_. Let *G* and *D* be the sets of genuine and predicted biclusters, respectively, then we call *ms(G, D)* and *ms(D, G)* the recovery and relevance score, respectively. *F*_*1*_-score is an overall measure and is defined to be 2 times the ratio between relevance × recovery and relevance + recovery.

### Programs Compared

We downloaded the R packages of ISA and FABIA, and the source code of BicPAMS, CPB and OPSM from their respective websites. We ran ISA and FABIA in R version 3.4.1, and compiled BicPAMS, CPB and OPSM on a linux workstation. We run the programs with the default settings, except for BicPAMS. We used sequential patterns gathered from multiple levels of expression {4, 7, 12, 20} and minimal columns {4, 5, 6}. For both FABIA and ISA, we set the “number of starting points” to be 1.1 times the number of hidden biclusters, and for QUBIC, UniBic and BicGO, we set “number of bicluters” to be the numbers of implanted biclusters.

### Generation of Artificial Datasets

We first produced background matrices using Gaussian distribution *N*(0, 1) with expectation 0 and standard deviation 1, and then implanted various trend-preserved biclusters in the background matrices. To implant an *s×t* bicluster in a background matrix A, we randomly selected an *s×t* submatrix in A and a row in the submatrix, and then permutated other rows in the submatrix such that the permutated rows are all trend-preserved with the selected row, resulting a trend-preserved bicluster. Clearly, the elements in the implanted bicluster have the same distribution as those in the background matrix. We also generate background matrices by implanting biclusters whose elements follow Gaussian distribution *N*(0, 1). To test the robustness of the algorithm, we added various levels of noises to the biclusters by adding to the elements in biclusters a residual *α* from *N*(0, *t*), where *t* was the noise level. The implanted biclusters were allowed to overlap with one another.

## Results

We tested BicGO by comparing it to seven other state-of-the-art tools, BicPAMS, CPB, FABIA, ISA, OPSM, QUBIC and UniBic on various artificial datasets as well as real datasets. Firstly, we tested their capability to identify both narrow and broad trend-preserved biclusters on artificial datasets. Secondly, we compared their robustness on artificial datasets with different noise levels. Finally, we applied them to real world gene expression datasets to test their ability to recover functionally related genes. Notice that the results in the experiments, see figures 1-3, were carried out on datasets without noises under consideration. Their robustness to noises will be compared in the robustness analysis section. Readers are referred to Supplementary Materials for results comparison on datasets with noises

**FIG. 1a.**
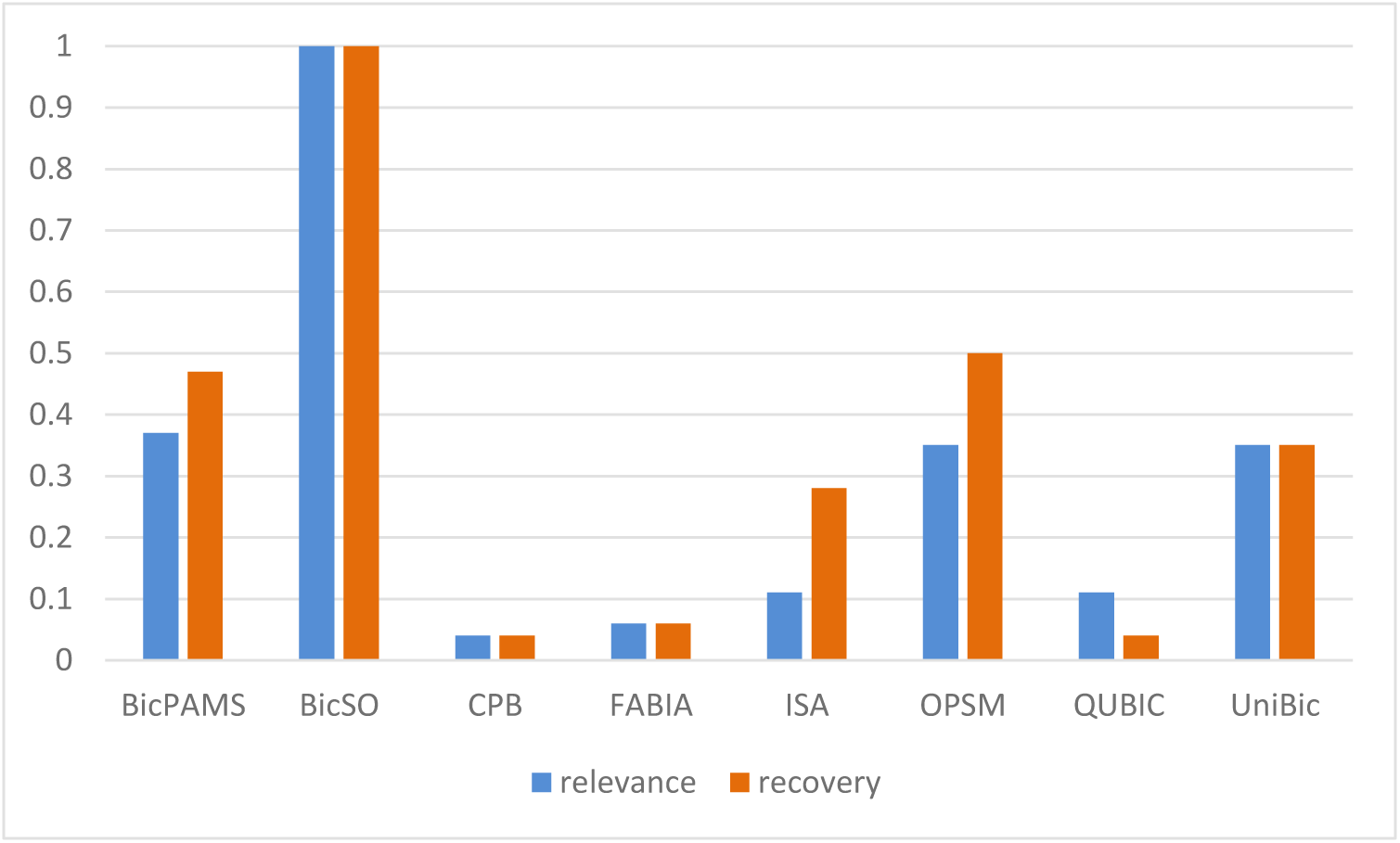
Comparisons in relevance and recovery scores achieved respectively by the eight salient tools on dataset in which three 100×10 trend-preserved biclusters were implanted, where the elements in implanted biclusters are distributed as same as those in background matrix which is generated from distribution N(0, 1).

**FIG. 1b.**
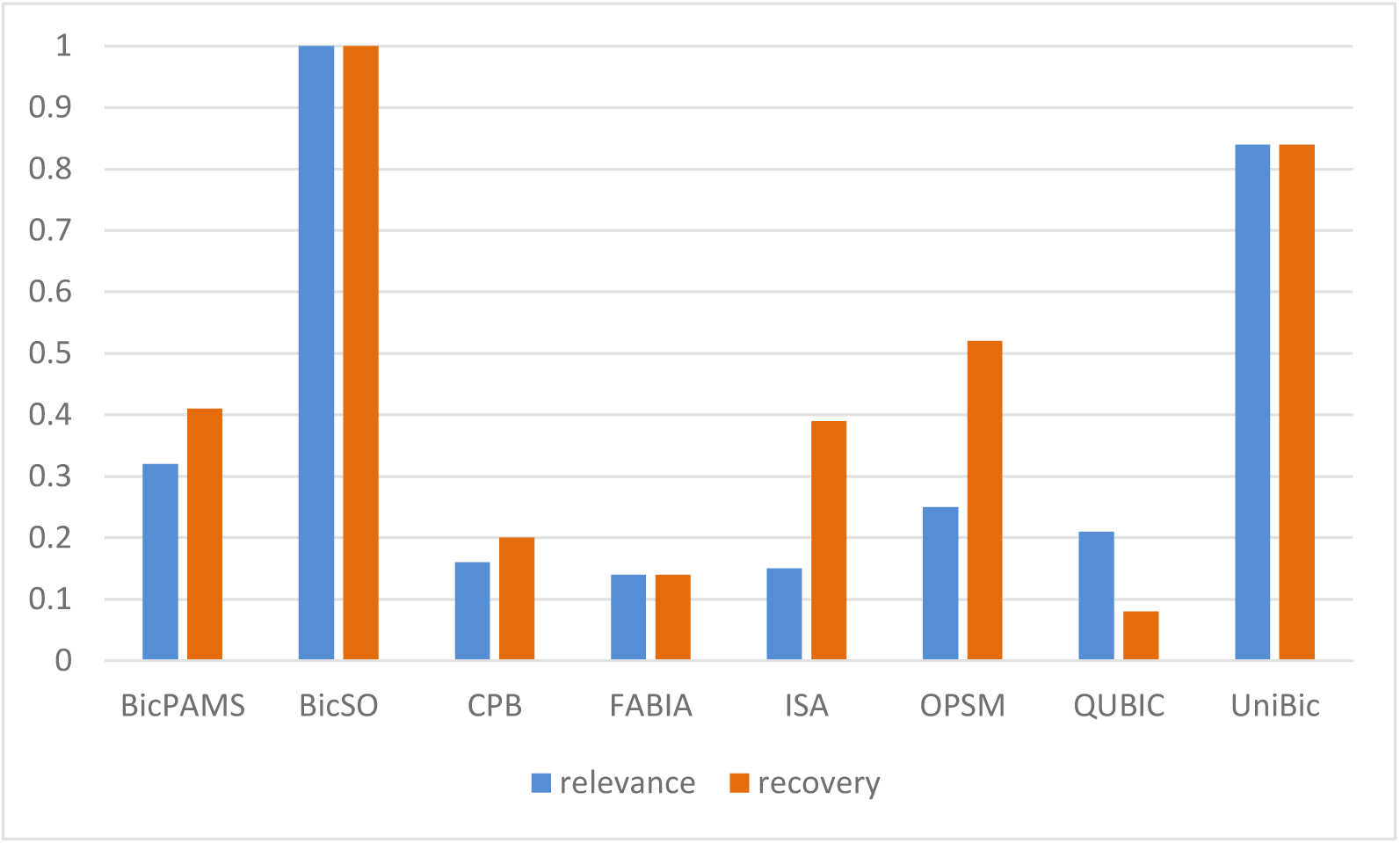
Comparisons in relevance and recovery scores achieved respectively by the eight salient tools on dataset in which three 100×20 trend-preserved biclusters were implanted, where the elements in implanted biclusters are distributed as same as those in background matrix which is generated from distribution N(0, 1).

**FIG. 1c.**
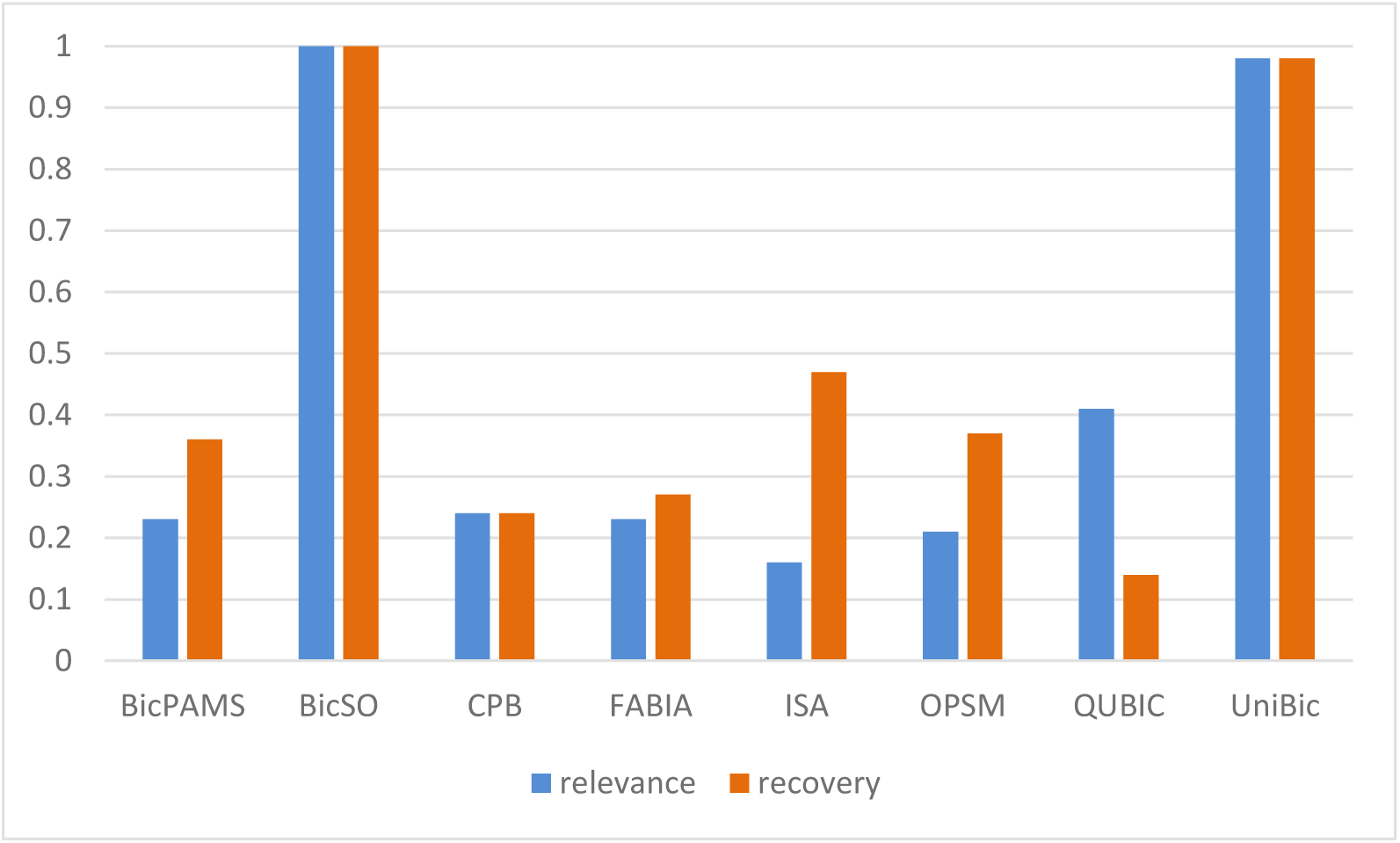
Comparisons in relevance and recovery scores achieved respectively by the eight salient tools on dataset in which three 100×30 trend-preserved biclusters were implanted, where the elements in implanted biclusters are distributed as same as those in background matrix which is generated from distribution N(0, 1).

### Test of BicGO on artificial datasets

#### Identification of narrow biclusters

In the real world, the biclusters to be detected can be quite narrow with many rows but only a few columns. We designed the column-based strategy tailored to identify relatively narrow biclusters while the data matrix might be arbitrary. To compare this strategy with the other programs for identification of narrow biclusters, we applied them to a 1000×100 background matrix implanted with trend-preserved biclusters of sizes 100×10, 100×20 and 100×30, respectively, where the elements in both background matrix and implanted biclusters obey the same distribution N (0, 1). For comparisons on datasets where the elements in implanted narrow biclusters have different distribution from those in background data, and moreover the implanted biclusters are of different types, e.g., const, shift, scale or shift-scale, the readers are referred to Supplementary Materials about narrow datasets. As shown in Fig. 1a, 1b and 1c, BicGO consistently achieved the highest relevance and recovery scores both are close to 1 in all the three datasets, followed by BicPAMS that is specifically tailored for trend-preserved narrower biclusters, but with much lower scores [(0.35, 0.47) in figure 1a, (0.32, 0.41) in figure 1b and (0.23, 0.36) in Fig. 1c]. The overall second-best performer, UniBic, increased from (0.35,0.35) to (0.98,0.98). There are two possible reasons why the other tools behave so poorly: (1) the implanted biclusters are too narrow to be detected as they were all developed based on row sequential pattern; and (2) the elements in both background matrix and implanted biclusters obey the same distribution while they were developed to find biclusters outstanding from the background noises. It is notably that all the tools but BicPAMS performed increasingly better with increase in the broadness of the biclusters to be identified, which is coincident with the fact that the broader biclusters are easier to be identified than are the narrow ones. It is worth noting that the column-based strategy worked equally well for identifying broader biclusters, e.g., its relevance and recovery scores were also close to (1, 1) in mining 100×30 biclusters (Fig. 1c).

#### Identification of broad biclusters

We designed the row-based strategy of BicGO aiming to improve UniBic on bad cases (see counterexample of UniBic in Supplementary Materials). We compared the row-based strategy with the other tools, but BicPAMS which was designed specifically for identification of narrow biclusters, on a 2000×200 background matrix generated from N(0, 1) with three shift-scale or trend-preserved biclusters implanted, whose rows and columns range from 50 to 70. The elements of the implanted biclusters follow either N(0, 1) or N(1, 1). The comparison results from figures 2a-d show that the row-based strategy consistently outperformed the other algorithms.

On the datasets with three 50 ×50, 60 × 60, 70 × 70 shift-scale biclusters implanted with their elements following N(0, 1) as same as followed by those in background matrix, the row-based strategy slightly outperformed UniBic with relevance and recovery scores (0.998, 0.998) vs (0.988, 0.988), but substantially outperformed all the other tools, with CPB holding the third place (0.800, 0.267) (Fig. 2a). On the datasets with three 50 ×50, 60 ×60, 70 ×70 trend-preserved biclusters implanted following the same distribution as the background data, the row-based strategy also slightly outperformed UniBic but substantially outperformed all the other tools (Fig. 2b). On the datasets with background data and implanted biclusters obeying N(0, 1) and N(1, 1) respectively, the row-based strategy substantially outperformed the other tools but slightly outperformed UniBic in both cases where the implanted biclusters were either shift-scale or trend-preserved (Fig. 2c, 2d).

**Fig. 2a.**
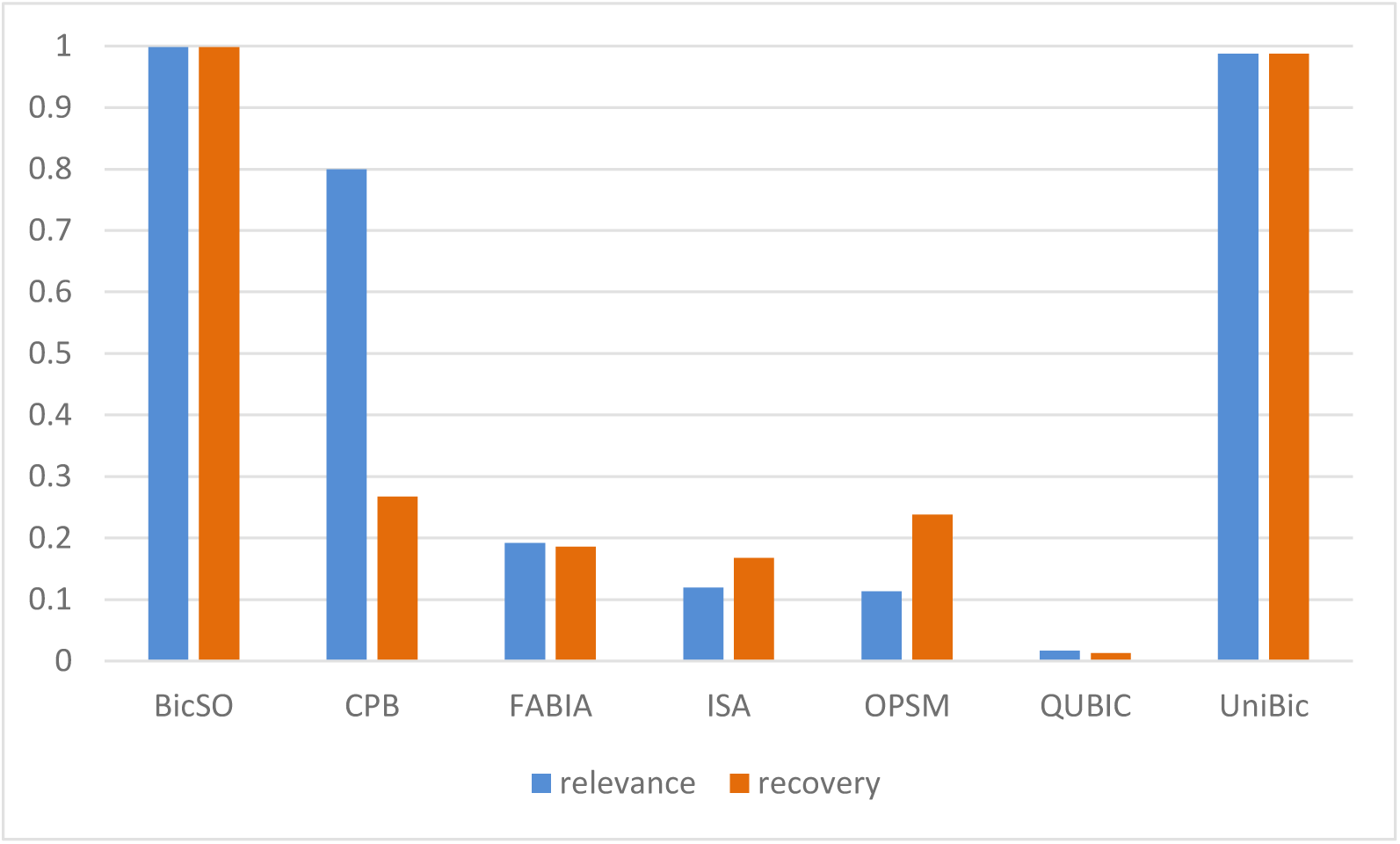
Comparisons in relevance and recovery scores achieved respectively by the seven salient tools on dataset in which three 50 ×50, 60 ×60, 70 ×70 shift-scale biclusters were implanted, where the elements in implanted biclusters are distributed as same as those in background matrix which is generated from distribution N(0, 1).

**FIG. 2b.**
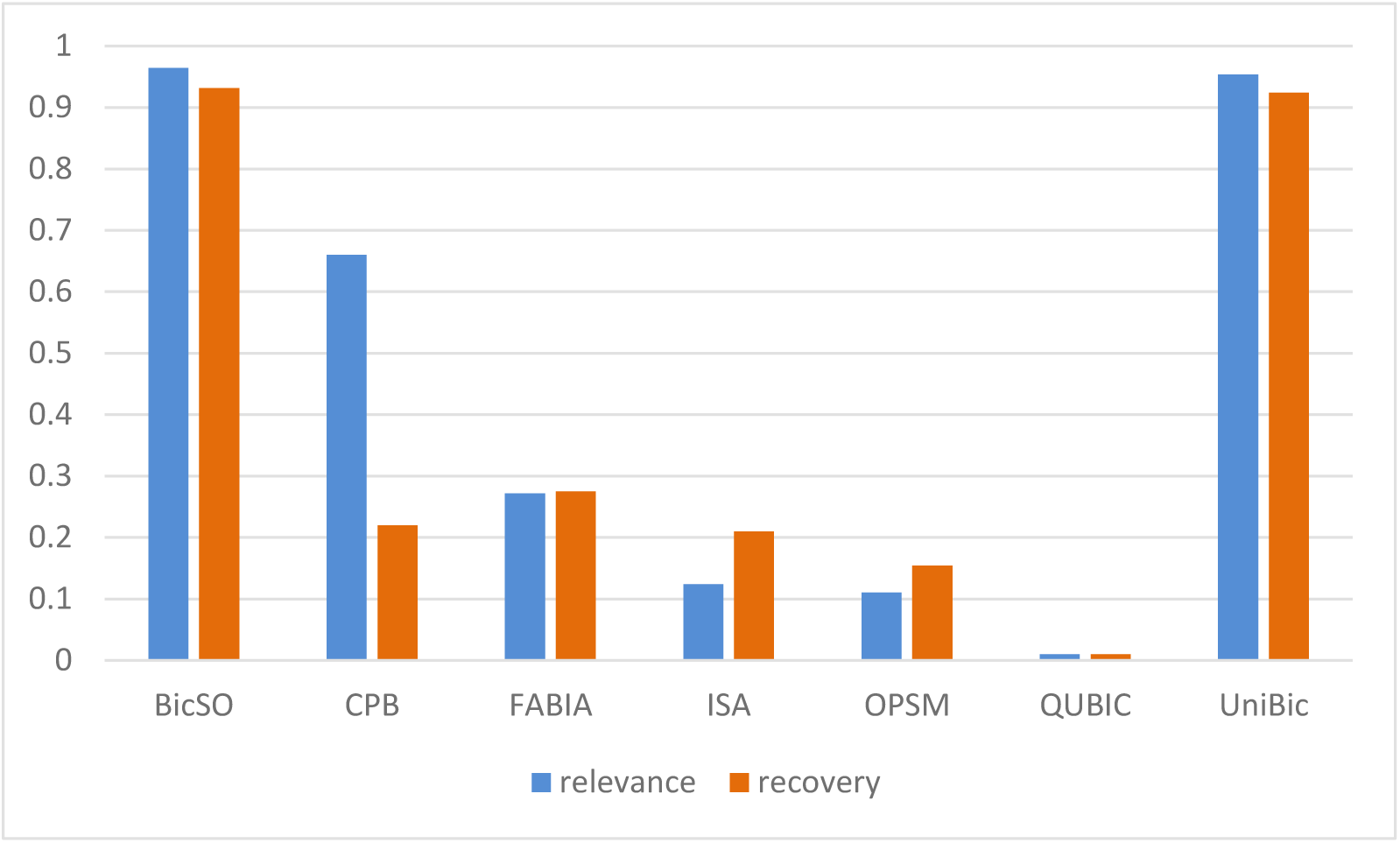
Comparisons in relevance and recovery scores achieved respectively by the seven salient tools on dataset in which three 50 × 50, 60 × 60, 70 × 70 trend-preserved biclusters were implanted, where the elements in implanted biclusters are distributed as same as those in background matrix which is generated from distribution N(0, 1).

**FIG. 2c.**
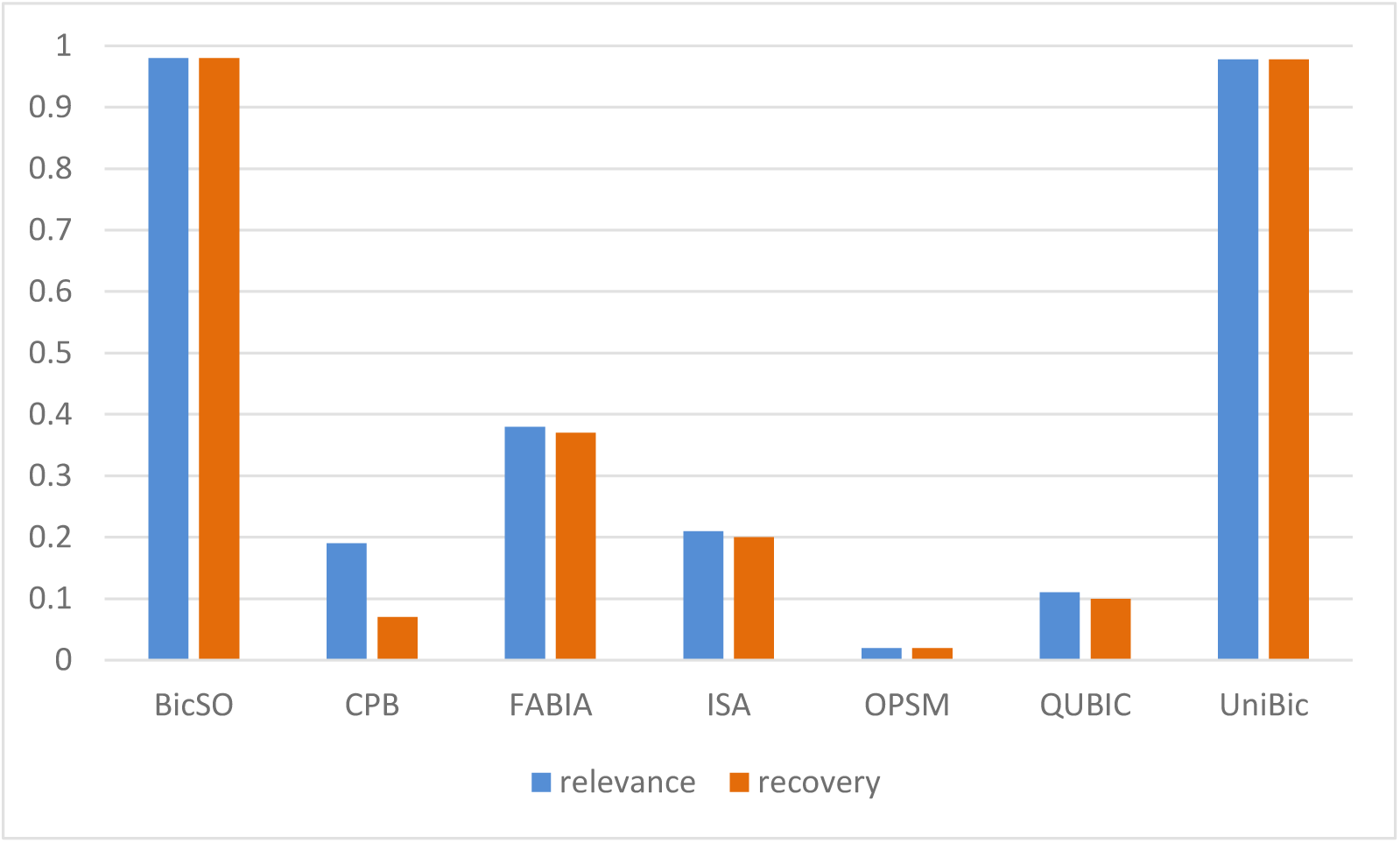
Comparisons in relevance and recovery scores achieved respectively by the seven salient tools on dataset in which three 50 × 50, 60 × 60, 70 × 70 shift-scale biclusters were implanted, where the elements in background matrix follow distribution N(0, 1), and those in implanted biclusters N(0, 1).

**FIG. 2d.**
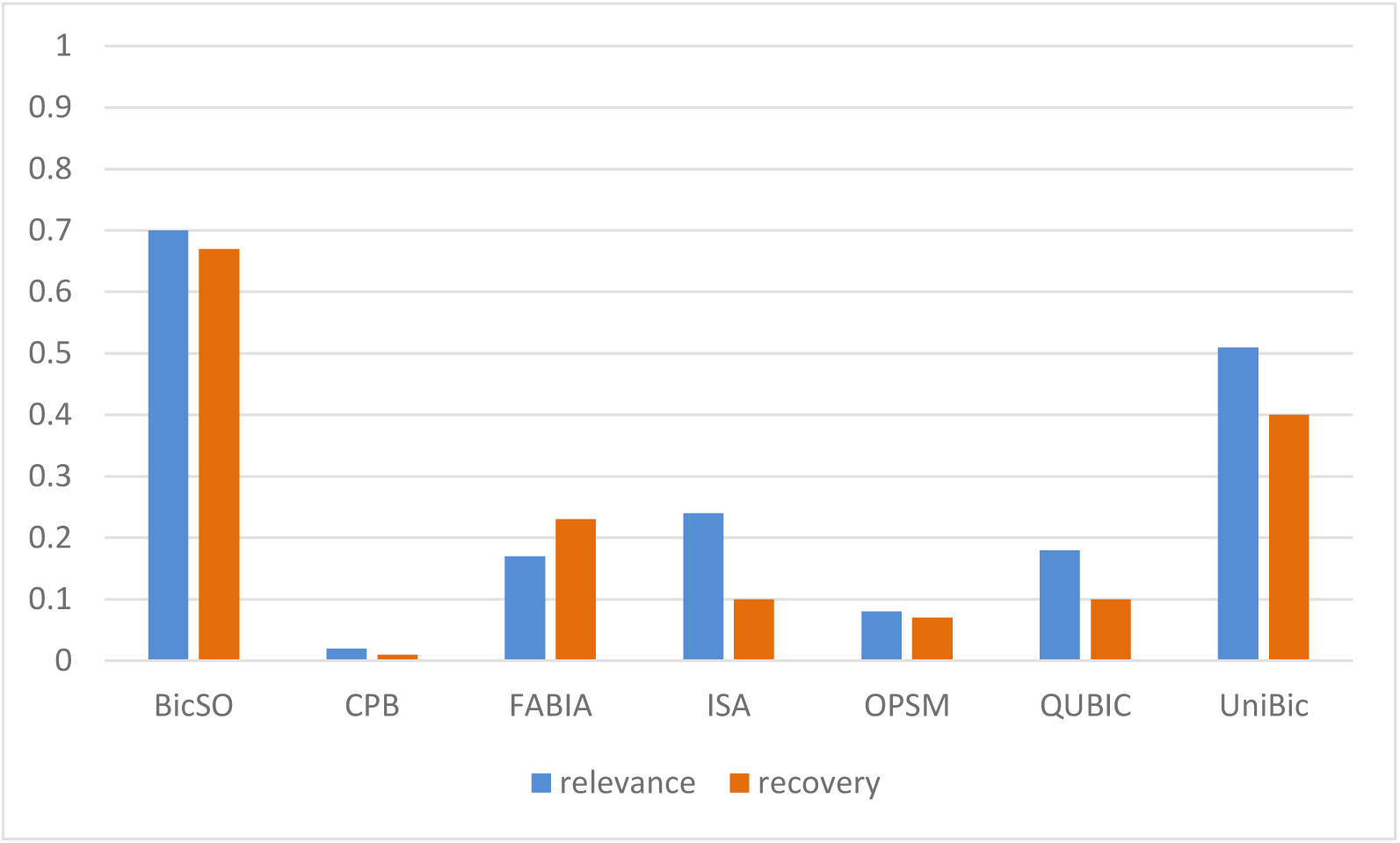
Comparisons in relevance and recovery scores achieved respectively by the seven salient tools on dataset in which three 50 × 50, 60 × 60, 70 × 70 trend-preserved biclusters were implanted, where the elements in background matrix follow distribution N(0, 1), and those in implanted biclusters N(0, 1).

#### Identification of both narrow and broad biclusters

In this subsection we tested all the eight salient tools on the dataset consisting of a 220×120 background matrix implanted with two broad biclusters of size 20×22 and two narrow ones of size 50 ×5, where the elements in both background matrix and implanted biclusters follow the same distribution N(0, 1). As shown in Fig. 3, BicGO reached the highest relevance and recovery scores (0.85, 0.84) vs (0.51, 0.47), followed by UniBic, demonstrating that BicGO had a remarkable advantage in detecting both narrow and broad biclusters owing to the two strategies of BicGO.

**FIG. 3:**
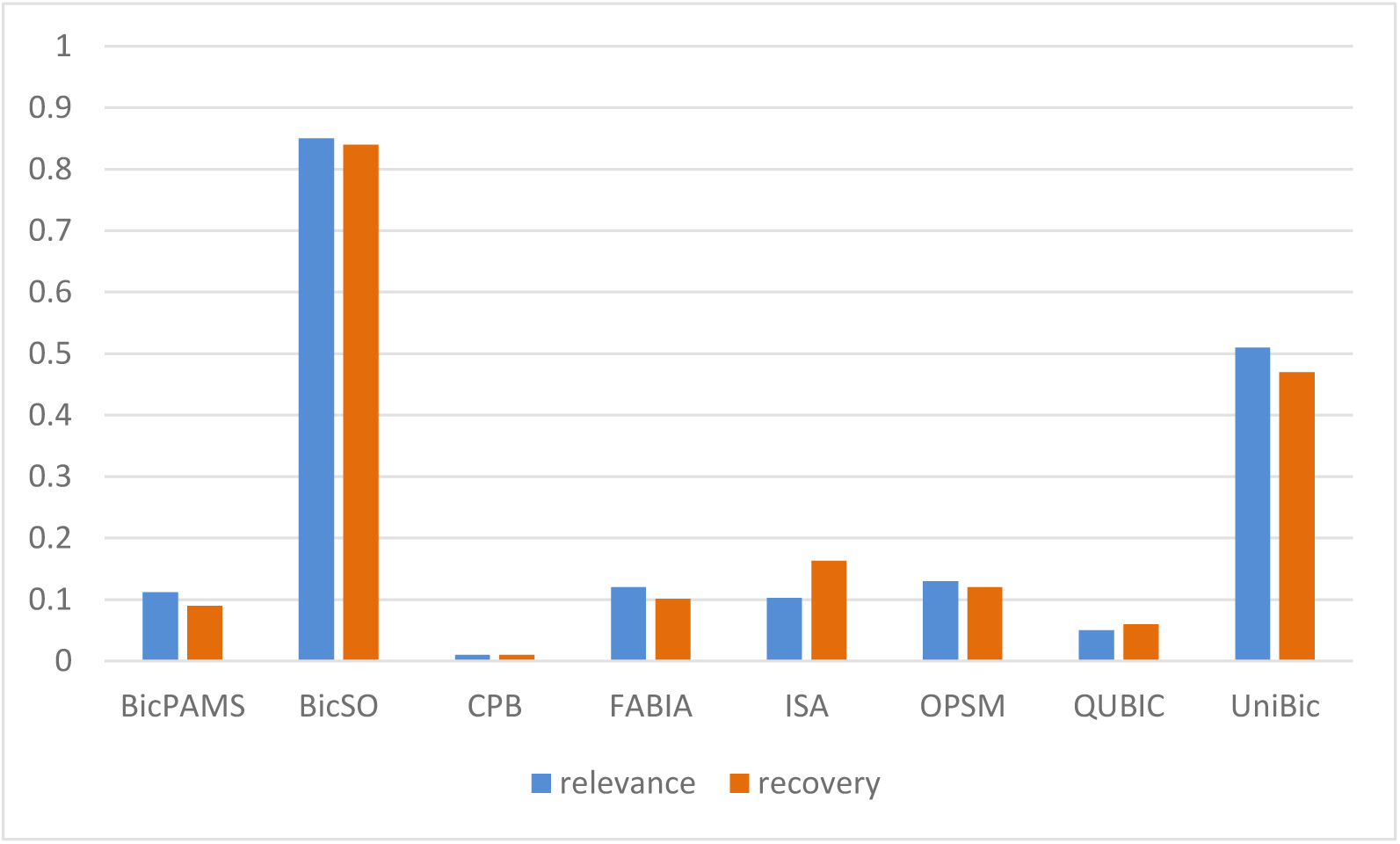
Comparisons in relevance and recovery scores achieved respectively by the eight tools on dataset in which both broad and narrow biclusters were implanted, where the elements in both background matrix and implanted biclusters follow the same distribution N(0, 1).

#### Robustness analysis

We next tested the impact of noises on the performance of the eight tools on a 500×30 background matrix in which a 100×10 bicluster implanted with variable noise levels. As shown in Fig. 4, BicGO consistently achieved the highest F_1_-scores on all the noise levels followed by UniBic or OPSM, both are significantly superior to the others across all the noise levels.

**FIG. 4.**
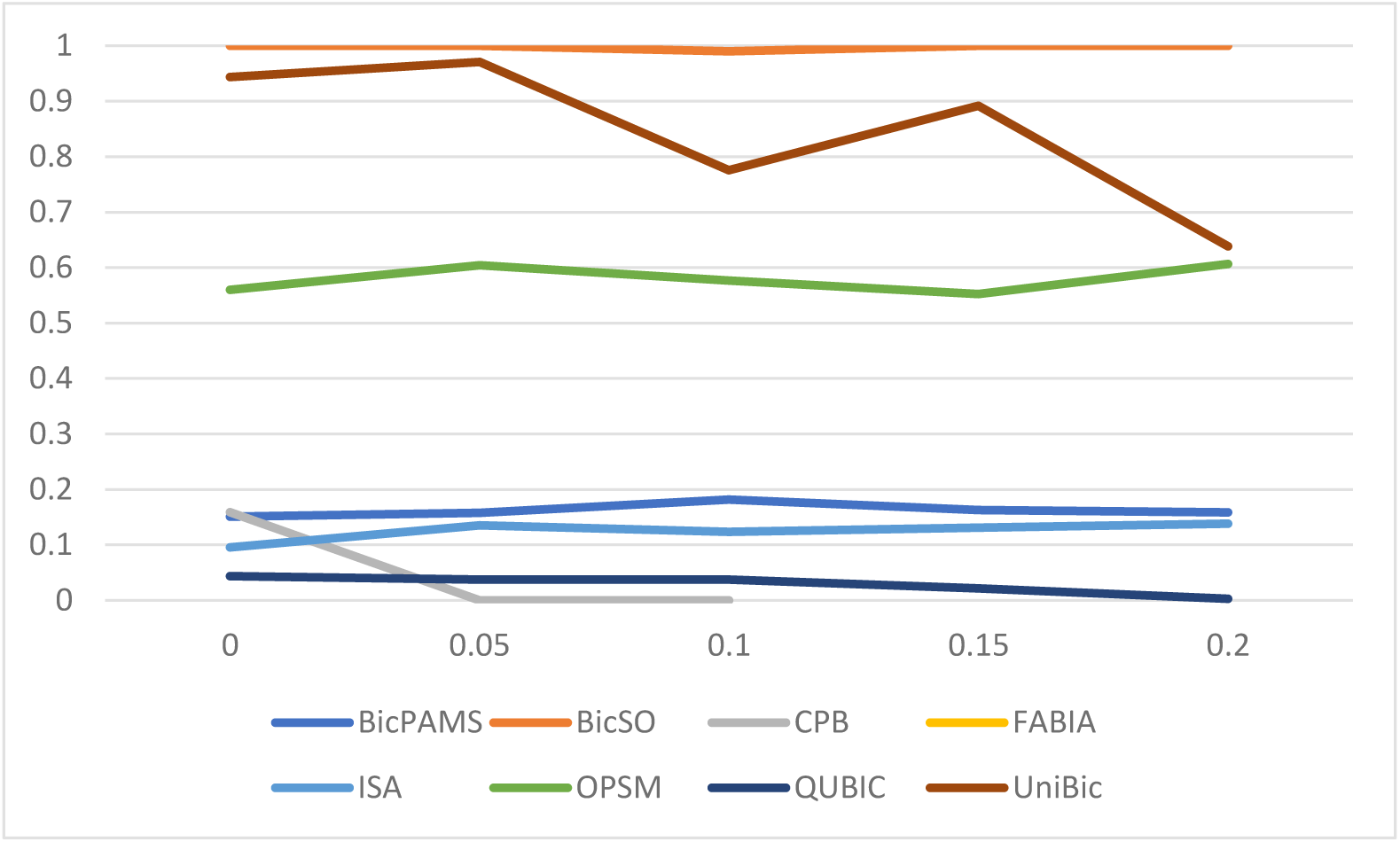
Comparison of distributions of F_1_-scores against noise levels achieved by the eight programs.

**FIG. 5.**
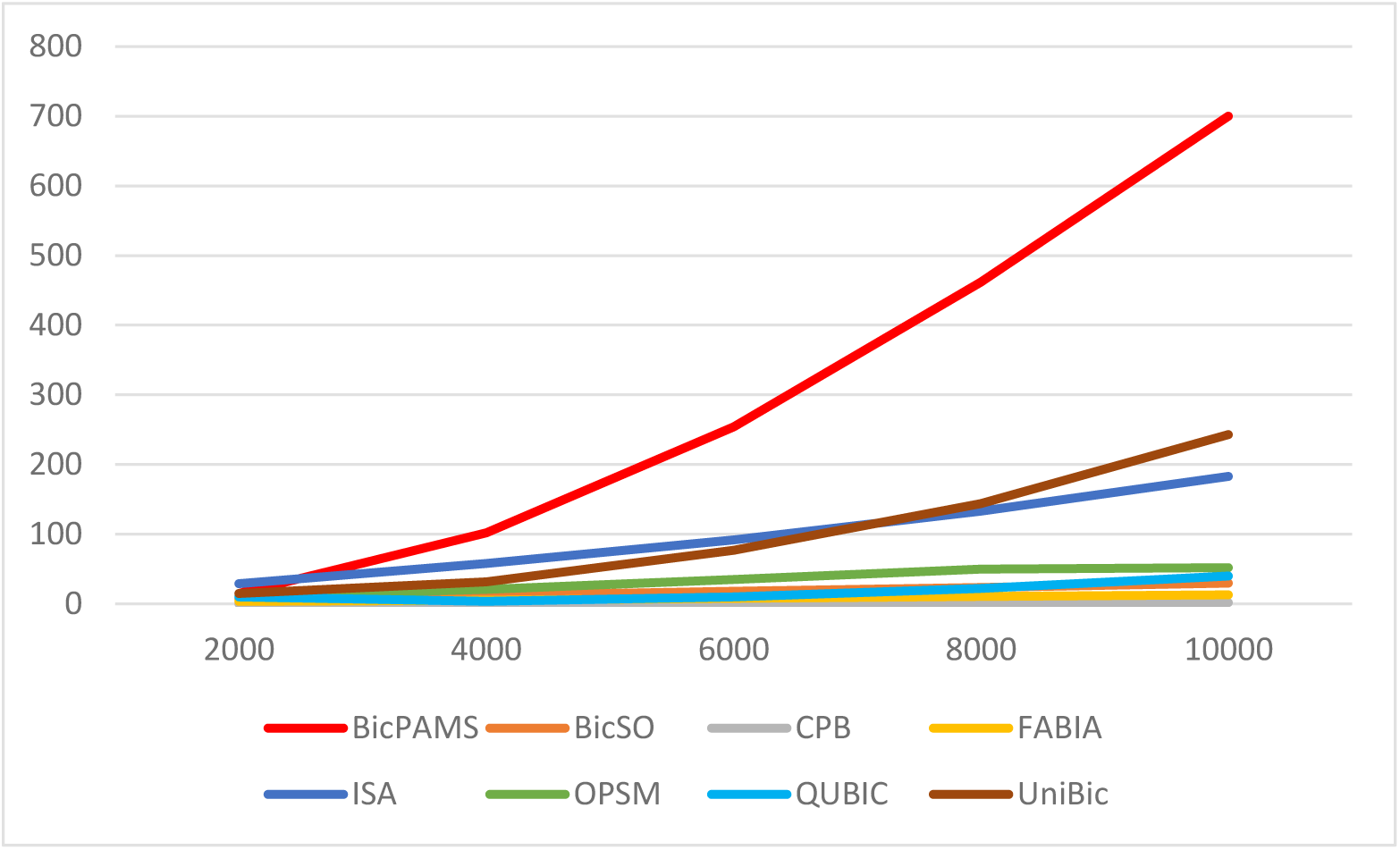
Comparisons of running time as a function of the number of rows in a data matrix with 50 columns.

### Performance of BicGO on real datasets

We further compared BicGO with the other seven algorithms on five real gene expression datasets from the GEO database: gds181, gds187, gds1406, gds2520 and gds3716 with more than 12,000 genes and dozens of conditions/samples (Table 1). We used the PCA method (ref.) to fix up the missing values. Since the true biclusters in the datasets are unknown, we used the GO terms enrichment to evaluate each bicluster discovered by each tool, with significant p-value 0.05. Since the eight tools may output different numbers of biclusters, we assessed their performances by the proportion of GO enriched biclusters. We ran BicGO by the column strategy with 50 biclusters output for each dataset. The results are summarized in Table 2. BicGO found a total of 163 significant enriched biclusters with the enriched rate of 65.2%, outperforming the other seven algorithms, followed by BicPAMS with an average enriched rate of 56.9%. QUBIC and CPB found more biclusters of 400 and 395, respectively, but achieved smaller enriched rates of 31.5% and 38.2%, respectively. ISA had the lowest enriched rate of 28% with 7 enriched biclusters among 24 biclusters found.

**Table 1.** Description of GDS datasets

| Datasets | Genes | Samples | Description |
| --- | --- | --- | --- |
| gds181 | 12626 | 84 | Large-scale analysis of the human Transcriptome |
| gds187 | 12487 | 12 | Progesterone and Hoxa-10 relationship |
| gds1406 | 12476 | 87 | Brain regions of various inbred strains |
| gds2520 | 12625 | 44 | Head and neck squamous carcinoma |
| gds3716 | 22283 | 42 | Breast cancer: Histologically normal breast epithelium |

**Table 2.** The results of GO enrichment analysis on eight GDS datasets

| Algorithm | Found | Enriched |
| --- | --- | --- |
| BicSO | 250 | 163 (65.2%) |
| BicPAMS | 187 | 109 (56.9%) |
| CPB | 395 | 151 (38.2%) |
| FABIA | 182 | 98 (53.8%) |
| ISA | 24 | 7 (28%) |
| OPSM | 75 | 34 (45%) |
| QUBIC | 400 | 126(31.5%) |
| UniBic | 314 | 149(47%) |

### Complexity estimation

Biclustering has been proved to be NP-hard even for the special case where the biclusters in the data matrix are constant. Therefore, a heuristic approach has to be taken to approximately identify biclusters in a data matrix in an effective and efficient way. Having demonstrated the effectiveness of BicGO compared to the seven existing salient tools, we estimate its efficiency. Given a A_m×n_ data matrix, where m is the number of rows and n the number of columns, let t be the biggest number of columns of the biclusters to be identified, o be the number of biclusters to be output (a parameter prespecified by users), and k be the smallest number of columns or rows of the biclusters to be identified. For the column-based strategy, BicGO takes O(n^2^m/k) comparisons to create a list of all the seeds, and O(kmn) comparisons to grow a seed to a bicluster. Therefore it takes O(n^2^m/k) + O(okmn) = O(max(n^2^/k, okn)m) comparison operations in total. For row-based strategy, BicGO takes O(n^2^(m/k)^2^k) to create a list of all the seeds, and O(kmn^2^) to grow a seed to a bicluster. Therefore it takes O(m^2^n^2^/k) + O(okmn^2^) = O(max{mn/k, okn}n^2^) in total. Figure 8 shows the running times on a laptop (with a CPU core-i7 6700hq and RAM 16G) of the algorithms on five background matrices with 50 columns and varying number (2000 ∼10000) of rows, each was implanted with a 100×10 bicluster. Clearly, BicGO using the column-based strategy is faster than most of the other programs.

## Discussion

Since Cheng and Church’s seminal work [1, 2], biclustering has been widely used in analyses of gene expression data, as it provides flexibility for identifying co-expressed genes under some but not necessarily all conditions, which the traditional clustering methods lack. However, most of the existing biclustering algorithms were designed to identify a rather special class of biclusters. In this paper, we developed a novel bicluster algorithm BicGO to discover trend-preserved biclusters of any shape no matter narrow or broad in any types of datasets. In both the column- and row-based strategies, BicGO first identifies globally optimized seeds, and then grows them into a full-sized bicluster using a greedy strategy. In the column-based strategy, we grow an optimum seed by greedily adding a new column into the current bicluster by repeatedly deciding whether or not a given numerical number belongs to a given closed interval. In the row-based strategy, we grow an optimum seed by greedily adding a new row to the current bicluster by finding the longest path in a pseudo-acyclic directed graph defined by two numerical vectors. Mathematically, both strategies globally optimize their solutions in every iteration. To our best knowledge, this is the first time for the biclustering algorithm to be built on global optimization in each iteration. When comparing BicGO with seven salient biclustering algorithms, we demonstrated that BicGO substantially and consistently outperforms all these algorithms in identifying various kinds biclusters implanted in simulated datasets as well as functionally enriched biclusters in real gene expression datasets. Therefore, BicGO will facilitate the analysis of gene expression data to elucidate transcriptional regulation networks.

## Acknowledgements

This work was supported by National Science Foundation of China (61432010, 61771009 to GL), and The National Key Research and Development Program of China (2016YFB0201702 to GL), NIH (R01GM106013 to ZS) and National Science Foundation (DBI1661332 to ZS). The funders had no role in study design, data collection and analysis, decision to publish, or preparation of the manuscript.

## Data Availability

Code, sample input data and usage instructions are available at the following Website: Code: https://www.dropbox.com/s/hsj3j96rekoks5n/BicGO.zip?dl=0

Data: https://www.dropbox.com/s/k6ydqr75wvsz29s/data.zip?dl=0

